# ESCRT-Independent Surface Receptor And Transporter Protein Degradation By The ILF Pathway

**DOI:** 10.1101/167411

**Authors:** Erin Kate McNally, Christopher Leonard Brett

**Author notes:** Correspondence: christopher.

## Abstract

Surface receptor and transporter protein down-regulation drives cell signaling, quality control and metabolism underlying diverse physiology. After endocytosis, proteins are delivered to endosomes where ESCRTs package them into intralumenal vesicles, which are degraded by acid hydrolases upon fusion with lysosomes. However, reports of ESCRT-independent surface protein degradation are emerging suggesting that alternative, non-canonical pathways exist. Using *Saccharomyces cerevisiae* as a model, here we show that in response to substrates, protein misfolding or TOR signaling, some internalized surface transporters (Hxt3, Itr1, Aqr1) bypass ESCRTs en route to the lysosome membrane where they are sorted into an area that is internalized as an intralumenal fragment (ILF) and degraded upon organelle fusion. This ILF pathway also degrades typical ESCRT client proteins (Mup1, Can1, Ste3) when ESCRT function is impaired. As the underlying machinery is conserved, we speculate that the ILF pathway is an important contributor to receptor and transporter down-regulation in all eukaryotes.

## HIGHLIGHTS and eTOC BLURB

- The ILF pathway degrades surface polytopic proteins that bypass ESCRTs
- Surface proteins are degraded by the ILF pathway in response to misfolding, TOR or substrates
- The ILF pathway compensates for ESCRT impairment
- The ILF pathway is important for receptor and transporter down-regulation

It is not clear how surface receptor and transporter down-regulation occurs independent of ESCRT function. Here, McNally and Brett show that internalized surface polytopic proteins that bypass ESCRTs are delivered to lysosome membranes where they are sorted and packaged for degradation by the intralumenal fragment (ILF) pathway upon homotypic organelle fusion.

## INTRODUCTION

Surface polytopic proteins including receptors, transporters and channels are internalized and sent to the lysosome for degradation (*Katzmann et al., 2002; Babst 2011; Henne et al., 2011*). Precise control of their surface levels underlies diverse physiology, including endocrine function, wound healing, tissue development, nutrient absorption and synaptic plasticity (*Katzmann et al., 2002; Palacios et al., 2005; Rodahl et al., 2008; Lobert et al., 2010; Zhou et al., 2010; Hislop and von Zastrow, 2011; Koumanov et al., 2012; Chassefeyre et al., 2015*). Damaged surface proteins are also cleared by this mechanism to prevent proteotoxicity (*Wang et al., 2011; Keener and Babst, 2013; Zhao et al., 2013*). To trigger this process, surface proteins are labeled with ubiquitin – in response to changing substrate levels, heat stress to induce protein misfolding or cellular signaling for example – and then selectively internalized by the process of endocytosis (*Blondel et al., 2004; Lewis and Pelham, 2009; MacGurn et al., 2011; Jones et al., 2012; MacDonald et al., 2012; Keener and Babst, 2013; Babst, 2014*). Within the cell, they are sent to endosomes where they encounter ESCRTs (Endosomal Sorting Complexes Required for Transport). These five protein complexes (ESCRT-0, ESCRT-I, ESCRT-II, ESCRT-III and the Vps4 complex) sort and package these internalized surface proteins into IntraLumenal Vesicles (ILVs; *Henne et al., 2011*). After many rounds, ILVs accumulate creating a mature MultiVesicular Body (MVB; *Nickerson et al., 2010; Hanson and Cashikar, 2012*). The MVB then fuses with lysosomes to expose protein laden ILVs to lumenal hydrolases for catabolism (*Katzmann et al., 2002*).

Although many examples of ESCRT-mediated protein degradation have been published (*see Babst, 2014*), reports of ESCRT-independent degradation of surface proteins are emerging (e.g. *Bowers et al., 2006; Theos et al., 2006; Leung et al., 2008; Silverman et al., 2013; Parkinson et al., 2015*). Furthermore, ILVs can be formed independent of ESCRT function and proteins recognized by ESCRTs continue to be degraded when ESCRTs are impaired (*Trajkovic et al., 2008; Blanc et al., 2009; Stuffers et al., 2009b; Edgar et al., 2014*). These realizations have led to one of the most prominent open questions in our field: What accounts for ESCRT-independent ILV formation and surface polytopic protein degradation?

Around the time when ESCRTs were discovered (*Katzmann et al., 2001*), Wickner, Merz and colleagues reported that an ILV-like structure called an IntraLumenal Fragment (ILF) is formed as a byproduct of homotypic lysosome fusion in the model organism *Saccharomyces cerevisiae* (*Wang et al., 2002*). Prior to lipid bilayer merger, fusogenic proteins and lipids concentrate within a ring at the vertex between apposing lysosomal membranes (*Wang et al., 2003; Fratti et al., 2004*). Upon SNARE-mediated membrane fusion at the vertex ring, the encircled area of membrane, called the boundary, is excised and internalized within the lumen of the fusion product where it encounters lysosomal hydrolases (*Mattie et al., 2017*).

We recently discovered that lysosomal polytopic proteins, e.g. ion and nutrient transporters, are selectively sorted into the boundary membrane for degradation in response to substrate levels, misfolding by heat stress or TOR (Target Of Rapamycin) signaling (*McNally et al., 2017*). Named the ILF pathway, this process functions independently of ESCRTs and instead relies on the fusion protein machinery for transporter sorting and ILF formation. Thus, this process performs similar functions as ESCRTs, except the mechanisms underlying protein sorting and packaging and their cellular locations are distinct. Thus, the possibility exists that surface polytopic proteins may also be degraded by the ILF pathway if they can be delivered to the lysosome membrane after internalization.

Can internalized surface proteins be delivered to lysosome membranes? To our knowledge, this hypothesis has not been formally tested. However, this proposition seems reasonable when considering the consequence of bypassing ESCRT function in the canonical MVB pathway (**Figure 1A**): In theory, any internalized surface polytopic protein that is not packaged into ILVs by ESCRTs (or returned to the surface) remains embedded within the outer membrane of the mature MVB. When this membrane merges with the lysosome membrane upon heterotypic fusion, these proteins are then exposed to the ILF machinery, which may package them for degradation. Thus, by simply avoiding recognition by ESCRTs, internalized surface proteins are––by default––delivered to lysosome membranes and the ILF pathway.

**Figure 1.**
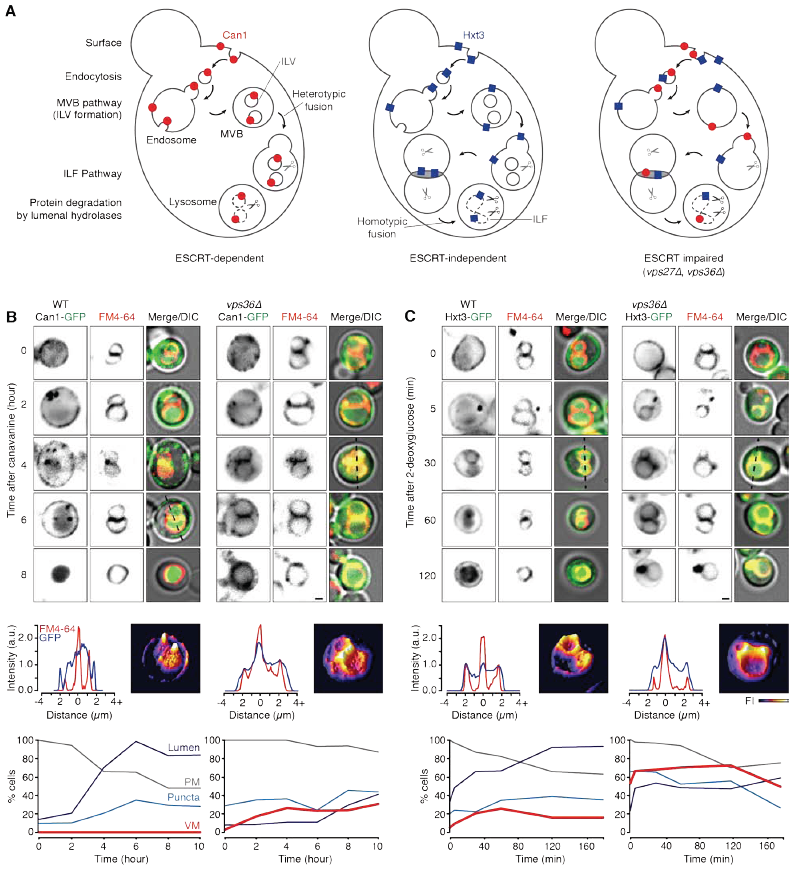
Internalized surface proteins take two routes to the lysosomal lumen for degradation. (**A**) Cartoon illustrating how surface membrane proteins can be sorted for degradation by the ESCRT or ILF pathways. (**B and C**) Fluorescence and DIC micrographs showing routes taken by Can1-GFP (**B**) or Hxt3-GFP (**C**) from the surface to the lysosome lumen in response to 37 µM canavanine or 200 µM 2-deoxyglucose over time in live wild type or *vps36*∆ cells treated with FM4-64 to label vacuole membranes. Scale bars, 1 µm. Middle panels include 3-dimensional GFP fluorescence intensity plots (right) and line plots (left) of GFP (blue) or FM4-64 (red) fluorescence intensity for micrographs where lines are indicated. Bottom panels indicate the proportion of the cell population that show GFP fluorescence on the plasma membrane (PM), intracellular puncta, vacuolar lysosome membrane (VM) or lysosomal lumen over time after treatment with a toxic substrate.

Upon reexamination of micrographs presented in earlier reports on receptor and transporter down-regulation, we found that some internalized surface polytopic proteins appear on lysosome membranes en route to the lumen for degradation, e.g. the high affinity tryptophan permease Tat2 (*Beck et al., 1999*), glucose transporters Hxt1 and Hxt3 (*O’Donnell et al., 2015*), peptide transporter Ptr2 (*Kawai et al., 2014*), and myo-inositol transporter Itr1 (*Nikko and Pelham, 2009*). We also noticed that most of these published studies do not directly assess whether ESCRTs are required for protein degradation. However, when the dependence on MVB formation was assessed, internalized surface proteins often accumulated on lysosome membranes when ESCRT function was disrupted, e.g. the general amino acid permease Gap1 (*Nikko et al., 2003*), ATP-binding cassette transporter Ste6 (*Krsmanovic et al., 2005*), and G-protein coupled receptor Ste3 (*Yeo et al., 2003; Shields et al., 2009; Prosser et al. 2010*). Thus, given that internalized surface transporters and receptors can appear on lysosome membranes, we decided to test the hypothesis that the ILF pathway represents an alternative, ESCRT-independent mechanism for degradation of surface polytopic proteins and may compensate for the loss of ESCRT function (**Figure 1A**).

## RESULTS

### Internalized surface proteins appear on lysosome membranes en route to the lumen for degradation

To test this hypothesis, we first confirmed that conventional ESCRT client proteins accumulate on vacuolar lysosome membranes when internalized into live cells that are missing components of the ESCRT machinery. To do so, we monitored cellular distribution of GFP-tagged transporters over time in populations of live *S. cerevisiae* cells using fluorescence microscopy and quantified their intracellular distribution (**Figure 1B**). GFP-tagged Can1, an arginine permease, is found on the plasma membrane prior to treatment with canavinine, a toxic arginine analog that triggers internalization and degradation of Can1 to prevent canavinine import and subsequent cell death (*MacGurn et al., 2011*). After treatment, Can1-GFP first appears on intracellular puncta and then later within the lysosomal lumen, but is never observed on the lysosome membrane.

However when VPS36 (a subunit of ESCRT-II; *Babst et al., 2002*) is deleted, Can1-GFP continues to be internalized after canavanine treatment where it accumulates on large punctae (reminiscent of vps class E compartments; *Raymond et al., 1992*) as well as on lysosome membranes, and eventually some fluorescence is observed in the lumen. We also found GFP-tagged Hxt3, a low-affinity, high capacity glucose transporter, accumulated on lysosome membranes en route to the lysosomal lumen after being internalized in response to 2-deoxyglucose, a toxic glucose analog (*O’Donnell et al., 2015;* **Figure 1C**). Importantly, this occurred in wild type cells and knocking out VPS36 did not prevent it from accumulating on lysosome membranes or within the lumen suggesting that Hxt3-GFP is processed for degradation by an ESCRT-independent mechanism. Because these surface polytopic proteins appear on the lysosome membrane after being internalized, we hypothesized that the ILF pathway may deliver them to the lysosome lumen for degradation.

### Surface transporters are sorted and packaged for degradation by the ILF pathway

To test this hypothesis, we first studied the lysosome membrane distribution of Hxt3-GFP, which does not seem to require ESCRTs for lysosomal degradation and is present in boundary membranes of docked lysosomes within live cells after treatment with 2-deoxyglucose (**Figures 1C and S1**). It is worth noting that the resident lysosomal polytopic protein Fet5-GFP was excluded from boundaries when cells were treated with 2-deoxyglucose (**Figure S1**), confirming that Hxt3-GFP sorting into the boundary is selective. Treating cells with heat stress to induce protein misfolding also caused Hxt3-GFP to appear in boundary membranes, similar to proteins that are degraded by the ILF pathway (**Figures 2A**; *McNally et al., 2017*). This discovery was not limited to Hxt3-GFP: we found that GFP-tagged Itr1 (a myo-inositol transporter; *Nikko and Pelham, 2009*) as well as Aqr1 (a major facilitator superfamily-type transporter that excretes amino acids; *Velasco et al., 2004*) take a similar route from the surface to the lysosomal lumen when live wild type cells were treated with heat stress (**Figure 2A**). Importantly, knocking out VPS36 did not affect sorting of these proteins into the boundary of docked lysosomes, confirming that they are sorted by an ESCRT-independent mechanism. We also found that Hxt3-GFP within the boundary was internalized during lysosome fusion events within live *vps36*∆ cells (**Figure 2B; Movie S1**), suggesting that ESCRT function is not necessary for protein delivery to the lumen for degradation.

**Figure 2.**
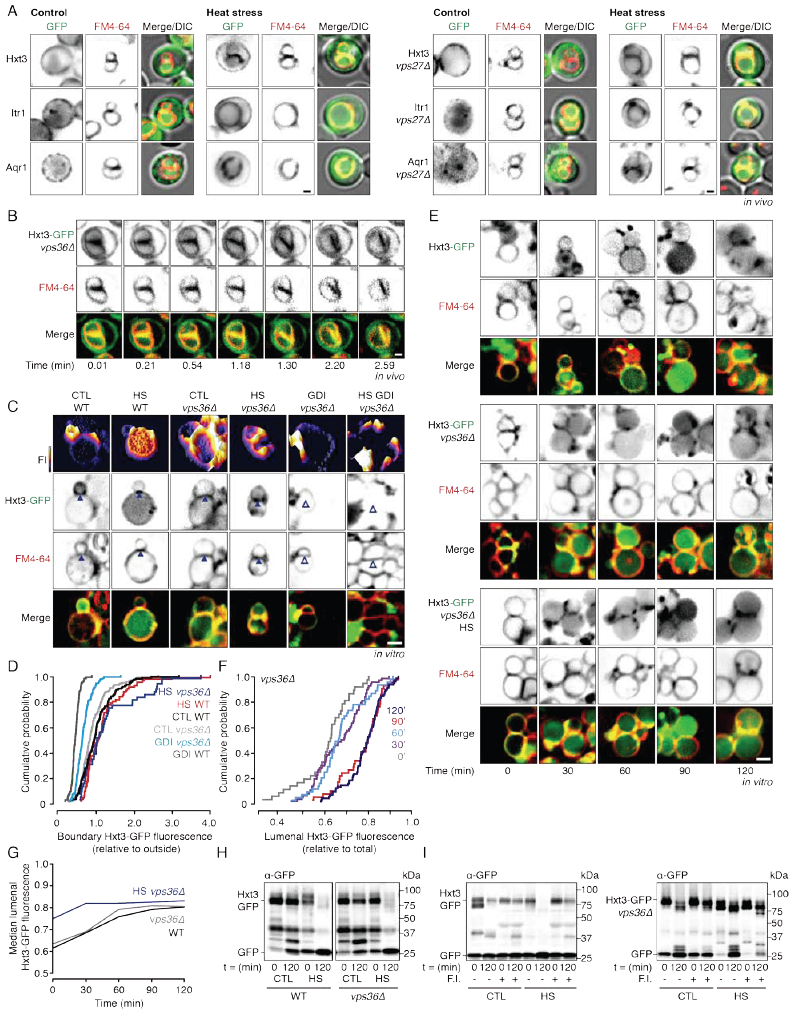
Some surface proteins are sorted for degradation by the ILF pathway. (**A**) Fluorescence and DIC micrographs of live wild type (left) or *vps27*∆ (right) cells expressing GFP-tagged Hxt3, Itr1 or Aqr1 before (control) and after heat stress for 15 minutes. (**B**) Images from time-lapse video showing homotypic lysosome fusion within live *vps36*∆ cells expressing Hxt3-GFP. See **Movie S1**. (**C**) Fluorescence micrographs of lysosomes isolated from wild type (WT) or *vps36*∆ cells after 30 minutes of fusion in the absence or presence of heat stress (HS) or rGdi1 and rGyp1-46 (GDI). 3-dimensional fluorescence intensity (FI) plots of Hxt3-GFP are shown. Arrowheads indicate Hxt3-GFP enrichment (closed) or exclusion (open) within the boundary membrane. Also see **Figure S2A**. (**D**) Cumulative probability plot of Hxt3-GFP fluorescence measured at boundaries between lysosomes isolated from wild type (WT) or *vps36*∆ cells after fusion in the presence or absence of heat stress (HS) or fusion inhbitors (rGdi1), as shown in (C; n ≥ 98). Also see **Figure S2B**. (**E**) Fluorescence micrographs of lysosomes isolated from wild type or *vps36*∆ cells expressing Hxt3-GFP acquired over the course of the fusion reaction in the absence or presence of heat stress (HS). (**F**) Cumulative probability plot of Hxt3-GFP fluorescence measured within the lumen of lysosomes isolated from *vps36*∆ cells after 0, 30, 60, 90 or 120 minutes of fusion, as shown in (E). Also see **Figure S2C**. (**G**) Median values of lumenal GFP fluorescence observed over time for three conditions studied (n ≥ 105). (**H and I**) Western blot analysis of Hxt3-GFP degradation before or after lysosomes isolated from wild type (WT) or *vps36*∆ cells underwent fusion for 120 minutes in the absence (CTL) or presence of heat stress (HS; H), or after fusion in the presence or absence of the fusion inhibitors (F.I.) rGdi1 and rGyp1-46 (I). Cells or isolated organelles were stained with FM4-64 to label vacuolar membranes. Scale bars, 1 µm in vivo, 2 µm in vitro.

Because the ILF machinery co-purifies with vacuolar lysosomes, we were able to develop cell-free assays to assess protein sorting, internalization and degradation by the ILF pathway (*McNally et al., 2017*). Thus, to further study how surface proteins may use this mechanism for degradation, we next isolated lysosomes from Hxt3-GFP expressing cells and imaged them 60 minutes after adding ATP to stimulate fusion *in vitro* (**Figure 2C**). Similar to findings made *in vivo*, Hxt3-GFP was sorted into the boundary during organelle fusion *in vitro*. Treating lysosomes with heat stress increased the amount of Hxt3-GFP sorted into the boundary (**Figure 2C and D**), suggesting that the machinery necessary to enhance Hxt3 degradation is present on lysosomes. Because the Rab GTPase Ypt7 is important for protein sorting into the ILF pathway (*McNally et al., 2017*), we next added the Rab inhibitors rGdi1 (a Rab-GTPase chaperone protein) and rGyp1-46 (the catalytic domain of the Rab GTPase activating protein Gyp1; *Brett et al., 2008*) to fusion reactions and found that Hxt3-GFP was no longer included in boundary membranes (**Figure 2C, 2D and S2**). Hxt3-GFP membrane distribution was not affected by deletion of VPS36, suggesting that sorting of Hxt3-GFP into the boundary requires Ypt7 and the fusion protein machinery, not ESCRTs.

To confirm that Hxt3 was internalized into the lumen upon lysosome fusion, we imaged lysosomes over the course of the fusion reaction *in vitro* (**Figure 2E**) and measured the GFP intensity within the lumen (**Figure 2F, 2G and S2**). As expected, Hxt3-GFP accumulated within the lumen over time after lysosome fusion was stimulated with ATP. Heat stress increased the rate and amount of lumenal GFP accumulation over time. Importantly, Hxt3-GFP internalization did not require VPS36, confirming that delivery to lumen did not require ESCRT function. Once delivered to the lumen of lysosomes, polytopic proteins embedded within ILFs are degraded by acid hydrolases (*McNally et al., 2017*). To assess proteolysis, we conducted western blot analysis to detect cleavage of GFP from Hxt3-GFP (**Figure 2H**) and found that more GFP was cleaved after lysosome fusion was stimulated *in vitro*. More Hxt3-GFP cleavage was further stimulated when lysosomes were treated with heat stress, consistent with sorting and internalization phenotypes observed. Furthermore, cleavage was unaffected by deleting VPS36 (**Figure 2H**) but was blocked by the fusion inhibitors rGdi1 and rGyp-46 (**Figure 2I**), confirming that the fusion machinery, not ESCRTs, was responsible for Hxt3-GFP degradation. Together, these findings indicate that in response to substrates or protein misfolding the surface transporter Hxt3-GFP is internalized and delivered to lysosome membranes where it utilizes the ILF pathway, and not ESCRTs, for degradation.

### ILF pathway degrades ESCRT client proteins when MVB formation is impaired

When ESCRT function is disrupted cells survive and surface proteins that are typically sorted and packaged by ESCRTs continue to be degraded although less efficiently (e.g. the manganese transporter Smf1; *Jensen et al., 2009*). In **Figure 1B** we show that Can1-GFP accumulates on lysosome membranes when VPS36 is deleted, suggesting that the ILF pathway may account for degradation. We confirmed that a similar distribution is observed for other ESCRT client proteins including GFP-tagged Ste3, a G-protein coupled receptor (*Shields et al., 2009*), and Mup1, a methionine permease (*MacDonald et al., 2012*; **Figure 3A**). Importantly, all three are present on boundary membranes formed between docked lysosomes within cells missing either VPS36 or VPS27 (a subunit of ESCRT-0; *Katzmann et al., 2003;* **Figure 3A and S3**), suggesting that disrupting function of ESCRT–0 or –1 results in the same outcome. As expected, all three surface proteins also accumulate in boundaries upon treatment with heat stress to induce protein misfolding (**Figure 3A**) when components of ESCRTs are deleted. Using Mup1-GFP as an example, we next monitored live *vps27*∆ cells and found that it is present within ILFs that form during organelle fusion (**Figure 3B; Movie S2**), suggesting that Mup1-GFP is sorted and packaged for degradation by the ILF pathway.

**Figure 3.**
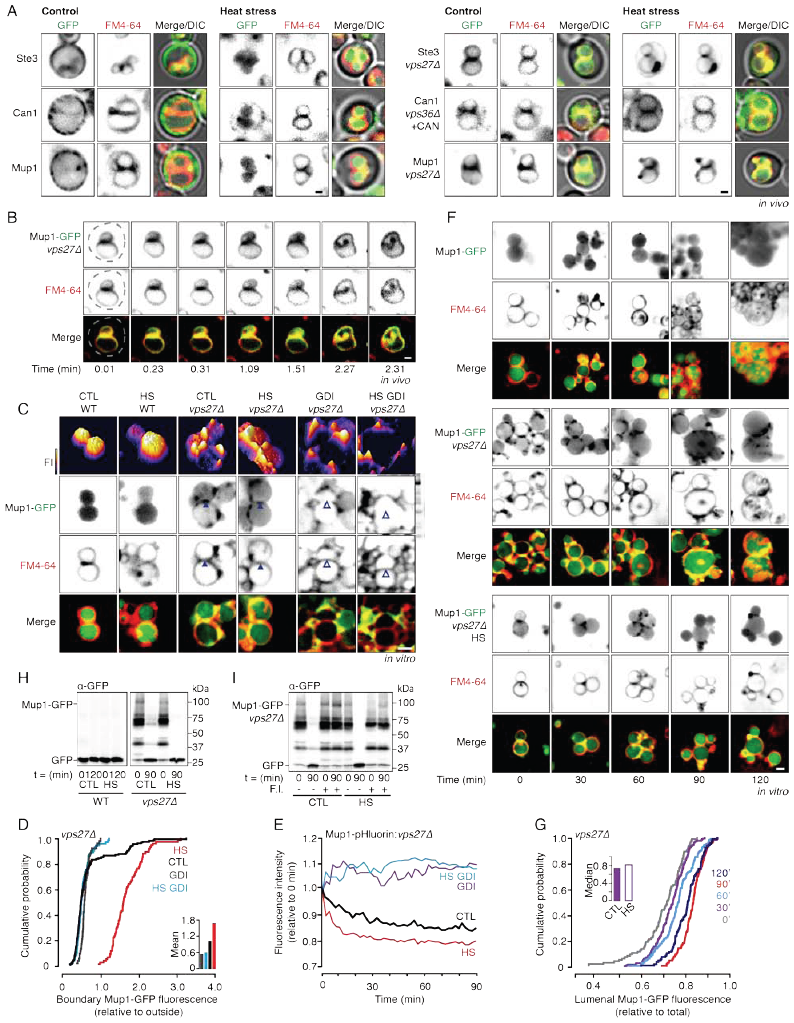
The ILF pathway compensates for ESCRTs when VPS27 is deleted. (**A**) Fluorescence and DIC micrographs of live wild type (left) and *vps27*∆ or *vps36*∆ (right) cells expressing GFP-tagged Ste3, Can1 or Mup1 before (control) and after heat stress for 30 minutes. Also see **Figure S3A**. (**B**) Images from time-lapse video showing homotypic lysosome fusion within live *vps27*∆ cells expressing Mup1-GFP. See Movie S2. (**C**) Fluorescence micrographs of lysosomes isolated from wild type (WT) or *vps36*∆ cells after 30 minutes of fusion in the absence or presence of heat stress (HS) or rGdi1 and rGyp1-46 (GDI). 3-dimensional fluorescence intensity (FI) plots of Mup1-GFP are shown. Arrowheads indicate Mup1-GFP enrichment (closed) or exclusion (open) within the boundary membrane. (**D**) Cumulative probability plot of Mup1-GFP fluorescence measured at boundaries between lysosomes isolated from *vps27*∆ cells after 30 minutes of fusion in the absence (CTL) or presence of heat stress (HS) or fusion inhibitors (GDI), as shown in (C), Inset shows mean values (n ≥ 115). (**E**) Fluorescence of lysosomes isolated from *vps27*∆ cells expressing Mup1-pHluorin during the *in vitro* fusion reaction under control conditions (CTL) or after heat stress (HS) in presence or absence of rGdi1 and rGyp1-46 (GDI). Also see **Figure S3B**. (**F**) Fluorescence micrographs of lysosomes isolated from wild type or *vps27*∆ cells expressing Mup1-GFP acquired over the course of the fusion reaction in the absence or presence of heat stress (HS). (**G**) Cumulative probability plot of Mup1-GFP fluorescence measured within the lumen of lysosomes isolated from *vps27*∆ cells after 0, 30, 60, 90 or 120 minutes of fusion, as shown in (**F**). Inset shows median values at 30 minutes for lysosomes from *vps27*∆ cells in the presence or absence of heat stress (n ≥ 118). Also see **Figure S3C**. (**H and I**) Western blot analysis of Hxt3-GFP degradation before or after lysosomes isolated from wild type (WT) or *vps36*∆ cells underwent fusion for 120 minutes in the absence (CTL) or presence of heat stress (HS; **H**), or after fusion in the presence or absence of the fusion inhibitors (F.I.) rGdi1 and rGyp1-46 (I). Cells or isolated organelles were stained with FM4-64 to label vacuolar membranes. Scale bars, 2 µm.

We confirmed this finding by isolating lysosomes from Mup1-GFP expressing cells and studying its membrane distribution during homotypic fusion *in vitro* (**Figure 3C**). Again, Mup1-GFP only appears on lysosomal membranes when VPS27 is deleted, where it is present in boundary membranes of docked lysosomes. Inducing protein misfolding with heat stress promotes sorting of Mup1-GFP into the boundary (**Figure 3C and D**), confirming that isolated lysosomes possess the molecular machinery required to process Mup1-GFP for degradation. Furthermore, addition of the Ypt7 inhibitors rGdi1 and rGyp1-46 prevents Mup1-GFP entry into the boundary, confirming that this process is dependent on the fusion machinery and not ESCRT function.

We next demonstrated that Mup1 was internalized into the lumen by tagging its cytoplasmic C-terminus with pHluorin, a pH-sensitive variant of GFP, and monitoring fluorescence over the course of the fusion reaction *in vitro* (**Figure 3E**). As expected, in the absence of VPS27, Mup1-pHluorin fluorescence decreases over the course of the fusion reaction, indicative of exposure to the acid lumen of the lysosome, in the presence or absence of heat stress. No change in Mup1-pHuorin was observed when lysosomes isolated from wild type cells were studied (Figure S3), as Mup1-GFP is exclusively found in the lumen (see **Figure 3C**). Addition of Ypt7 inhibitors prevented internalization (**Figure 3E**) confirming that the fusion machinery was required for Mup1-pHluorin internalization. To confirm this result, we imaged isolated lysosomes at different time points over the course of the fusion reaction in vitro (**Figure 3F**) and measured GFP intensity within the lumen (**Figures 3G and S3**). As Mup1-GFP is exclusively found within the lumen of lysosomes isolated from wild type cells, lumenal GFP intensity does not change over time. However, in the absence of VPS27, Mup1-GFP is initially present on lysosome membranes and accumulates within the lumen over time. Heat stress increases lumenal Mup1-GFP, consistent with an increase of Mup1-GFP observed in the boundary (**Figure 3D**) and increased Mup1-pHluorin internalization upon organelle fusion (**Figure 3E**).

Once inside the lumen, Mup1-GFP should be degraded by acid hydrolases. To confirm, we conducted western blot analysis to detect cleavage of GFP from Mup1 before or 120 minutes after the homotypic lysosome fusion was stimulated with ATP *in vitro* (**Figure 3H**). Organelles isolated from wild type cells only contained cleaved GFP, confirming that Mup1-GFP was being sent to lysosomes for degradation prior to isolation. However, intact Mup1-GFP is detected on lysosome preparations from *vps27*∆ cells and is cleaved after fusion is stimulated. As predicted, treating lysosomes isolated from *vps27*∆ cells with heat stress increases GFP cleavage (**Figure 3H**) and the fusion inhibitors rGdi1 and rGyp1-46 blocks GFP cleavage (**Figure 3I**), consistent with effects on Mup1-GFP sorting and internalization (**Figure 3C–G**). Together these findings indicate that when ESCRT function is impaired, internalized surface polytopic proteins that are typically degraded by the ESCRT pathway are instead shunted to lysosome membranes where they are sorted and packaged for degradation by the ILF pathway.

### The ILF pathway degrades surface proteins in response to TOR signaling

Thus far we have shown that the ILF pathway degrades internalized surface transporters and receptors in response to toxic substrates for cell survival, or in response to protein misfolding for cellular quality control. Another important stimulus of protein degradation is TOR signaling (*Laplante and Sabatini, 2012*): TOR kinase is thought to be activated by low cytoplasmic amino acid levels or ribosome inactivity to promote degradation of soluble cytosolic proteins by the proteasome (*Rousseau and Bertolotti, 2016*), surface polytopic proteins by ESCRTs (*MacGurn et al., 2011*) and lysosomal polytopic proteins by the ILF pathway (*McNally et al., 2017*), which in turn raises free amino acid levels and closes a negative feedback loop (*Settembre et al., 2013*). Given the newfound role of the ILF pathway in surface polytopic protein degradation, we next sought to determine if this process also responds to TOR using cycloheximide (CHX) to trigger TOR signaling (see *MacGurn et al., 2011; McNally et al., 2017*). As predicted, we found that surface Hxt3-GFP was internalized and appeared on membranes and in the lumen of lysosomes in live cells in response to CHX, in the presence or absence of VPS36 (**Figure 4A**), suggesting that CHX-mediated down-regulation of Hxt3-GFP does not require ESCRTs. Similar observations were made for Mup1-GFP in cells lacking VPS27 (**Figure 4A**) as well as other ESCRT client proteins (**Figure S4**), suggesting that CHX triggers degradation of these surface proteins by the ILF pathway when ESCRT function is impaired. To confirm these observations, we recorded lysosome fusion events within live *vps27*∆ or wild type cells and observed boundary-localized Mup1-GFP or Hxt3-GFP, respectively, being internalized into the lumen upon fusion (**Figure 4B; Movies S3 and S4**). Together, these results suggest that TOR signaling also induces surface protein degradation by the ILF pathway.

**Figure 4.**
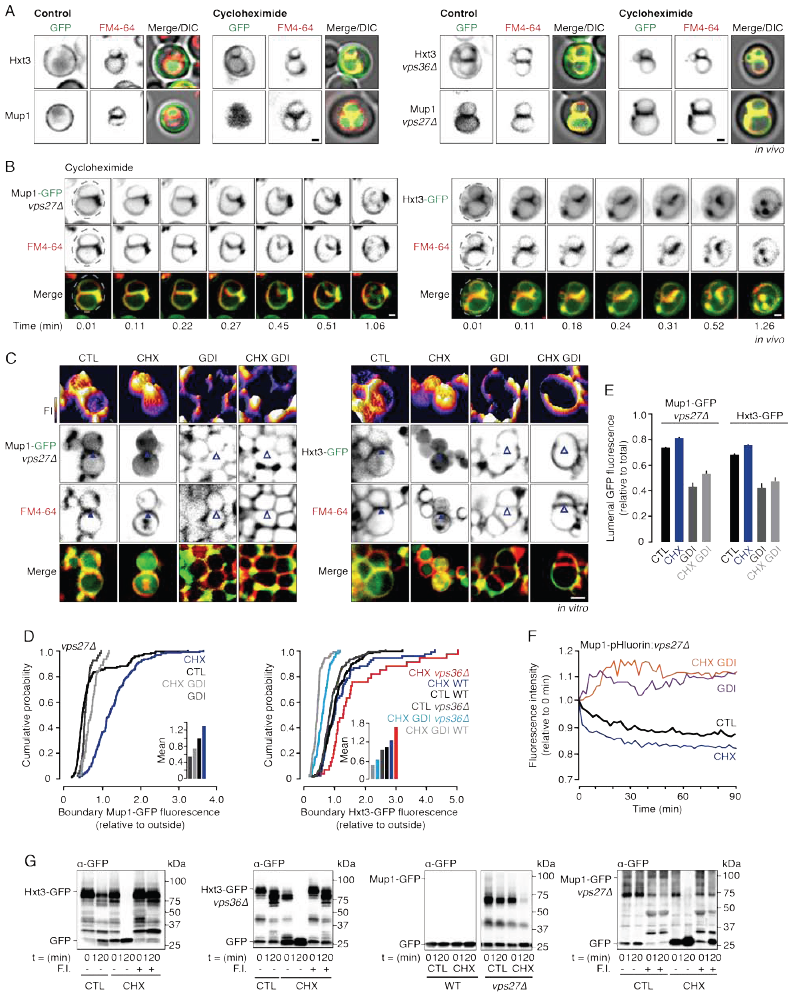
The ILF pathway degrades surface proteins in response to TOR activation. (**A**) Fluorescence and DIC micrographs of live wild type (left) or ESCRT impaired (*vps27*∆ or vps36∆; right) cells expressing GFP-tagged Hxt3 or Mup1 before (control) and after treatment with 100 µM cycloheximide (CHX) for 30 minutes. Also see **Figure S4A**. (**B**) Images from time-lapse video showing homotypic lysosome fusion within live *vps27*∆ cells expressing Mup1-GFP (see Movie S3) or wild type cells expressing Hxt3-GFP (see **Movie S4**) after CHX treatment. (**C**) Fluorescence micrographs of lysosomes isolated from wild type (WT) or *vps27*∆ cells after 30 minutes of fusion in the absence or presence of CHX or rGdi1 and rGyp1-46 (GDI). 3-dimensional fluorescence intensity (FI) plots of Mup1-GFP or Hxt3-GFP are shown. Arrowheads indicate GFP enrichment (closed) or exclusion (open) within the boundary membrane. Also see **Figure S4B and C**. (**D and E**) Cumulative probability plots of Mup1-GFP or Hxt3-GFP fluorescence measured at boundaries between (**D**) or within the lumen of (**E**) lysosomes isolated from wild type (WT) or *vps27∆* cells after fusion in the presence or absence of CHX or fusion inhibitors (GDI), as shown in (**C**; n ≥ 110). (**F**) Fluorescence of lysosomes isolated from *vps27*∆ cells expressing Mup1-pHluorin during the *in vitro* fusion reaction under control conditions (CTL) or after cycloheximide treatment (CHX) in presence or absence of rGdi1 and rGyp1-46 (GDI). Also see **Figure S4D**. (**G**) Western blot analysis of Mup1-GFP or Hxt3-GFP degradation before or after lysosomes isolated from wild type (WT) or ESCRT impaired (*vps27*∆ or *vps36∆*) cells underwent fusion for 120 minutes in the absence (CTL) or presence of CHX, or after fusion in the presence or absence of the fusion inhbitors (F.I.) rGdi1 and rGyp1-46. Cells or isolated organelles were stained with FM4-64 to label vacuolar membranes. Scale bars, 1 µm in vivo, 2 µm in vitro.

We next confirmed these findings in vitro using cell-free assays to further validate our hypothesis (**Figures 4C and S4**). We find that in the absence of VPS27, Mup1-GFP is enriched in boundary membranes upon treatment with CHX in vitro (**Figure 4D**), confirming that the TOR machinery that responds to CHX co-purifies with isolated lysosomes (*McNally et al., 2017*). We made similar findings for Hxt3-GFP in the absence or presence of VPS36 (**Figure 4D**), confirming that ESCRTs function was not important for Hxt3-GFP sorting in response to CHX. By measuring the lumenal GFP fluorescence intensity, we found that CHX caused more Hxt3-GFP and Mup1-GFP to accumulate inside lysosomes after membrane fusion was stimulated for 60 minutes in vitro (**Figure 4E**). We verified this finding by monitoring Mup1-pHluorin fluorescence over the course of the lysosome fusion reaction in vitro (**Figure 4F**), confirming that CHX stimulated protein internalization during fusion. We next confirmed that CHX also triggered more Hxt3-GFP and Mup1-GFP protein degradation by examining GFP cleavage by Western blot analysis (**Figure 4G**). Importantly, the lysosome membrane fusion inhibitors rGdi1 and rGyp1-46 blocked Hxt3-GFP and Mup1-GFP protein sorting (**Figure 4C and D**), internalization (**Figure 4E and F**) and degradation (**Figure 4G**). Furthermore, these events occurred in the absence of key components of the ESCRT machinery (VPS27 or VPS36; **Figures 4C–G and S4**). Thus, we conclude that the ILF pathway is responsible for degradation of some surface proteins (e.g. Hxt3-GFP) and degrades others (Mup1-GFP) to compensate for ESCRT impairment in response to TOR activation by CHX.

## DISCUSSION

### The ILF pathway degrades surface proteins without ESCRTs

Here we demonstrate that some internalized surface polytopic proteins are delivered to the vacuolar lysosome membrane where they are sorted into ILFs for degradation upon organelle fusion independent of ESCRT function (**Figures 1C, 2 and 4**). Previous reports of surface protein degradation often only include micrographs indicating the surface distribution of a protein before internalization, and its presence in the lumen of lysosomes afterwards (e.g. *Berkower et al., 1994; Egner et al., 1995; Shields et al., 2009; Keener and Babst, 2013; Zhao et al., 2013; Ghaddar et al., 2014; O’Donnell et al., 2015*); and many important papers originally describing degradation of surface receptors or transporters do not include any micrographs (e.g. *Hicke and Reizman, 1996*). Because the outcomes of both ESCRT and ILF pathways are equal, evidence showing the intracellular route of luminal delivery now seems important. For the ESCRT pathway, this requires imaging fixed and stained samples by electron microscopy to visualize proteins on small ILVs within endosomes (*Haigler et al., 1979; Futter et al., 1996; Klumperman and Raposo, 2014*) as current methods of light microscopy cannot distinguish whether proteins are present on ILVs or limiting membranes of endosomes.

However for the ILF pathway, here we show surface transporters being delivered to the lumen during homotypic lysosome fusion in real time within live cells using HILO microscopy (e.g. **Figures 2B and 4B, Videos S1 and S4**). These include sugar transporter Hxt3, as well as the peptide transporter Aqr1 and myo-inositol transporter Itr1, but re-examination of micrographs in previous reports reveal the presence of other internalized surface transporters on lysosome membranes as well (e.g. Tat2; *Beck et al., 1999*), suggesting that ILF-dependent degradation of surface proteins may be widespread in *S. cerevisiae*. These transporters and the underlying fusion machinery responsible for the ILF pathway (*Nickerson et al., 2009; Spang, 2016*) are evolutionarily conserved, suggesting that a similar process may regulate surface glucose (Hxt/GLUT gene family) or myo-inositol (Itr/HMIT) transporters levels important for cellular metabolism or nutrient (re-) absorption in human epithelial cells within the ileum or kidney for example (*Smith et al., 2013; Chen et al., 2015; Schneider, 2015*). Given that the mechanism of degradation has only been elucidated for only a handful of an estimated ~5,500 polytopic proteins encoded by the human genome (*Fagerberg et al., 2010*), we speculate that the ILF pathway could be equally important as the ESCRT pathway for determining surface protein lifetimes and thus may underlie diverse physiology.

We also show that the ILF pathway sorts and packages internalized surface proteins that are typically processed for degradation by ESCRTs when MVB formation is impaired (**Figures 1B, 3 and 4**). Unlike protein sorting into the ESCRT pathway, we were able to visualize their internalization by the ILF pathway in real time within live cells (**Figures 3B and 4B; Videos S2 and S3**). These include the ESCRT client proteins Mup1, Ste3 and Can1 that have been extensively used to characterize this pathway in *S. cerevisiae* (e.g. *Lin et al., 2008*). This may not be limited to internalized surface proteins, as biosynthetic cargo such as lysosomal acid hydrolases (e.g. the carboxypeptidase Cps1 and polyphosphatase Phm5) also utilize the MVB pathway for delivery to the lysosomal lumen, and deleting components of ESCRTs redistributes it on the lysosome membrane (a classic phenotype used to discover and characterize the ESCRT pathway; e.g. *McNatt et al., 2007*). Thus, we speculate that compensation by the ILF pathway may mediate residual surface protein degradation previously observed when ESCRTs are impaired (*Jensen et al., 2009*), thus permitting cell survival when the primary process for sorting and packaging membrane proteins for degradation is dysfunctional. Loss-of-function mutations in human orthologs of ESCRT components are linked to cancers and neurodegenerative disorders for example and etiology is thought to involve improper cargo degradation (*Saksena and Emr, 2009; Stuffers et al., 2009a; Alfred and Viccari, 2016*). Thus, assuming the ILF pathway also functions in human cells, it is tempting to speculate that up-regulating the ILF pathway by increasing lysosome numbers and activity through TFEB (Transcription Factor EB) signaling for example (*Ferguson, 2015; Napolitano and Ballabio, 2016*), may offer a therapeutic strategy to treat ESCRT-related diseases.

### Coordination of cellular protein degradation pathways for survival and physiology

What determines whether a surface protein uses the ESCRT or ILF pathways for degradation? Here we show that protein sorting into the ILF pathway is triggered by the same stimuli that mediate entry into the ESCRT pathway: changes in substrate levels (e.g. *Nikko and Pelham, 2009;* **Figure 1**), protein misfolding by heat stress (*Keener and Babst, 2013;* **Figures 2 and 3**), or TOR signaling (*MacGurn et al., 2011; Jones et al., 2012;* **Figure 4**). In the ESCRT pathway, this is mediated by cargo protein recognition by adapter proteins (e.g. arrestins; *Lin et al., 2008; Nikko and Pelham, 2009*) and ubiquitylation by E3-ligases (e.g. Rsp5; *MacDonald et al., 2012*) at surface or endosomal membranes. Although the mechanism responsible for protein sorting by the ILF pathway has not been elucidated, we have preliminary evidence suggesting that adapter proteins and E3-ligases are required for protein soring into the ILF pathway (*data not shown*), suggesting that both pathways recognize ubiquitylated proteins, which my explain why the ILF pathway can degrade proteins that are typically substrates for the ESCRT pathway. Furthermore, the same labeling machinery is present at the plasma membrane, endosomes and lysosomal membranes (e.g. *Li et al., 2015a; Li et al., 2015b*), suggesting that proteins are ubiquitylated in a similar manner at these sites and thus different patterns of protein ubiquitylation are unlikely. The presence of similar labeling machinery on the plasma and lysosome membranes may also explain why more Mup1-GFP is sorted into the ILF pathway and degraded when isolated lysosomes are treated with CHX or heat stress in vitro (**Figures 3 and 4**).

The underlying mechanism may also be ubiquitin-independent. This is because ubiquitylation is not required for sorting of all protein cargoes into the ESCRT pathway: some proteins bind to chaperones such as Sna3 to mediate degradation (*Regiorri and Pelham, 2001; McNatt et al., 2007; MacDonald et al., 2012*). Sna3 has a paralog called Sna4 that is exclusively found on lysosome membranes (*Pokyrzwa et al., 2009*) and is sorted into the ILF pathway (*McNally et al., 2017*). Thus, we hypothesize that perhaps chaperone binding specificity may determine which pathway is used by surface polytopic proteins for degradation. For example, Hxt3 may preferentially interact with Sna4 instead of Sna3. If so, internalized Hxt3 avoids Sna3-mediated incorporation into ILVs by ESCRTs at the endosome and remains on the limiting membrane. Upon MVB-lysosome fusion, Hxt3 is delivered to lysosome membranes where it can encounter Sna4 and be degraded by the ILF pathway. We are currently testing this hypothesis and anticipated results will help reveal unique and shared mechanisms that underlie these different cellular protein degradation pathways, giving us a better understanding of how they orchestrate global changes in protein turn over that occur during cell differentiation, the cell cycle or cellular aging for example.

### The ILF pathway as an alternative for other ESCRT-related cellular physiology

Could the ILF pathway also be responsible for ESCRT-independent ILV formation? Like ESCRTs, the ILF machinery selectively sorts and packages proteins into what are essentially large intralumenal vesicles. Thus, we speculate that homotypic fusion likely contributes to generating intralumenal vesicles observed within lysosomes that are needed for proper lipid catabolism (*Kallunki et al., 2013*). However, the underlying machinery is also responsible for heterotypic lysosome membrane fusion events between autophagosomes (*Jiang et al., 2014*) and MVBs (*Luzio et al., 2009; Epp et al., 2011*). Moreover, paralogous machinery underlies endosome membrane fusion required for earlier trafficking events during endocytosis (*Balderhaar and Ungermann, 2013; Spang, 2016; Chou et al., 2016*). Thus we speculate that ILF formation during organelle fusion may occur at other sites in the cell, including during homo- or heterotypic endosome fusion, which may account for ILVs within endosomes observed when ESCRTs are depleted (*Trajkovic et al., 2008; Blanc et al., 2009; Stuffers et al., 2009b; Edgar et al., 2014*) or for observed ILVs that are much larger than those derived by ESCRTs (e.g. *Fairn et al., 2011*). Although incredibly speculative, this ESCRT-independent mechanism may also account for exosome formation when lysosomes or lysosome-related organelles containing ILVs fuse with the plasma membrane (*Reddy et al., 2001; Jaiswal et al., 2002; Blott and Griffiths, 2002*). New methods of super-resolution fluorescence microscopy or electron microscopy are necessary to test this hypothesis as these organelles, ILVs and exosomes are expected to be smaller (e.g. 25-50 nm in diameter; see *Palay and Palade, 1955; Hanson and Cashikar, 2012; Edgar et al., 2014*) than those observed during homotypic vacuolar lysosome fusion (up to 1 µm in diameter based on boundary membrane lengths observed; *Mattie et al., 2017; McNally et al., 2017*). If true, these findings will confirm that the ILF pathway is as important as the ESCRT pathway for cellular physiology.

## EXPERIMENTAL PROCEDURES

### Yeast strains and reagents

All *Saccharomyces cerevisiae* strains used in this study are listed in **Table S1**. Biochemical and yeast growth reagents were purchased from either Sigma-Aldrich, Invitrogen or BioShop Canada Inc. Proteins used include recombinant Gdi1 purified from bacterial cells using a calmodulin-binding peptide intein fusion system (*Brett and Merz, 2008*) and recombinant Gyp1-46 (the catalytic domain of the Rab-GTPase activating protein Gyp1) purified as previously described (*Eitzen et al. 2000*). Reagents used in fusion reactions were prepared in 20 mM Pipes-KOH, pH 6.8, and 200 mM sorbitol (Pipes Sorbitol buffer, PS).

### Fluorescence microscopy

Images were acquired with a Nikon Eclipse TiE inverted microscope equipped with a motorized laser TIRF illumination unit, Photometrics Evolve 512 EM-CCD camera, CFI ApoTIRF 1.49 NA 100x objective lens, and 488 nm or 561 nm (50 mW) solid-state lasers operated with Nikon Elements software. Cross sectional images were recorded 1 μm into the sample and resulting micrographs and processed using ImageJ software (National Institutes of Health) and Adobe Photoshop CC. Images shown were adjusted for brightness and contrast, inverted and sharpened with an unsharp masking filter. Fluorescence intensity profiles of GFP fluorescence were generated using ImageJ software.

### Live cell microscopy

Live yeast cells were stained with FM4-64 to label vacuole membranes and prepared for imaging using a pulse-chase method as previously described (*Brett et al., 2008*). For examining vacuolar localization of plasma membrane proteins, cells were incubated at 37 °C for 15 minutes for heat stress (**Figure 2A**) after FM4-64 staining. For other stress treatments yeast cells were incubated at 37°C for 30 minutes (**Figure 3A**) and incubated with 100 μM cycloheximide for 90 minutes at 30 °C (**Figure 4A**). Time-lapse videos were acquired at 30°C using a Chamlide TC-N incubator (Live Cell Instruments) with cells plated on coverslips coated with concavalin-A (1 mg/ml in 50 mM HEPES, pH 7.5, 20 mM calcium acetate, 1 mM MnSO_4_). Yeast cells expressing Mup1-GFP were back diluted in Synthetic Complete (SC) media lacking methionine (Takara Bio USA, Inc.) for live cell microscopy. To assess surface protein degradation in response to substrate addition, yeast cells expressing Can1-GFP or Hxt3-GFP wild type and when VPS36 is deleted were stained with FM4-64 for one hour at 30 °C in SC media. After two washes, cells were resuspended in fresh SC media with addition of either 37 μM canavinine (Sigma-Aldrich) for 2, 4, 6 or 8 hours (Can1) or with 500 μM 2-Deoxy-D-glucose (Sigma-Aldrich) for 5, 30, 60 or 120 minutes (Hxt3 and Fet5). After incubation, cells were washed and resuspended in SC media prior to imaging.

### Vacuole isolation and homotypic vacuole fusion

Vacuoles were isolated from GFP derivative yeast cells as previously described (*Haas, 1995*) and fusion reactions were prepared using 6 μg of isolated vacuoles in standard fusion reaction buffer with 0.125 M KCl, 5 mM MgCl_2_, 1 mM ATP and 10 μM CoA. Vacuolar membranes were stained with FM4-64 by treating vacuoles with 3 μM FM4-64 for 10 minutes at 27 °C. Reactions were incubated at 27 °C for 60 minutes, unless otherwise noted, and placed on ice prior to visualization by microscopy. Where indicated, vacuoles were incubated in the absence or presence of 3.2 μM Gyp1-46 and 4 μ M rGdi. For heat stress treatment, vacuoles were pretreated for 5 minutes at 37 °C before addition to the fusion reaction and incubation for 30 minutes at 27°C. Where indicated, vacuoles were pretreated with 100 μM cycloheximide for 15 minutes at 27°C and incubated for an additional 15 minutes with the fusion reaction.

### pHluorin-based assay to detect protein internalization

Ecleptic pHluorin was cytoplasmically tagged to lysosomal membrane proteins (see *Prosser et al., 2010*) and reactions were prepared as previously described (*McNally et al., 2017*). Data shown are representative traces with values normalized to time zero, n≥4.

### Western blot analysis

Samples were prepared from isolated vacuoles from yeast strains expressing a GFP-tagged vacuolar membrane protein as previously described (*McNally et al., 2017*) and probed for α-GFP (Abcam, Cambridge, UK). Samples were repeated a minimum of three times. Gels were imaged using GE Amersham Imager 600 by chemiluminescence and edited and prepared using Adobe Photoshop and Illustrator CC software.

### Data analysis and presentation

Cumulative probability measurements were calculated using isolated vacuole micrographs of docked vacuoles only. GFP fluorescence intensities were measured using ROI 4x4 pixels in diameter and acquiring a fluorescence value on the outer membrane, within the lumen and on the boundary membrane of docked vacuoles using ImageJ software. For assessing GFP fluorescence in the boundary membrane, micrographs taken 60 minutes into the fusion reaction were assessed. For lumenal GFP fluorescence, micrographs were assessed at 0 (ice), 30, 60, 90 and 120 minutes into the fusion reaction and correlated to individual vacuole size (boundary length, surface area and circumference). Data are reported as mean ± SEM. Comparisons were calculated using Student two-tailed *t*-test, *P* values < 0.05 indicate significant differences.

Acquired data was plotted using Synergy KaleidaGraph 4.0 software and figures were prepared using Adobe Illustrator CC software.

## AUTHOR CONTRIBUTIONS

C.L.B and E.K.M. conceived the project. E.K.M. performed experiments and prepared all data for publication. E.K.M. and C.L.B. wrote the paper.

## ACKNOWLEDGEMENTS

We thank the Canadian Foundation for Innovation and Natural Sciences and Engineering Research Council of Canada for generous support of the Centre for Microscopy and Cellular Imaging at Concordia University. This work was supported by NSERC grant RGPIN/403537-2011 to C.L.B.

## SUPPLEMENTARY INFORMATION

**Figure S1 related to Figure 1.**
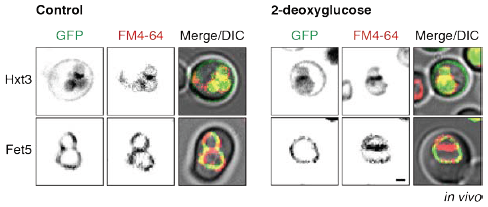
Fluorescence and DIC micrographs of live wild type cells expressing GFP-tagged Hxt3 or Fet5 before (control) and after addition of 2-deoxyglucose for 30 minutes. Scale bar, 1 µm.

**Figure S2 related to Figure 2.**
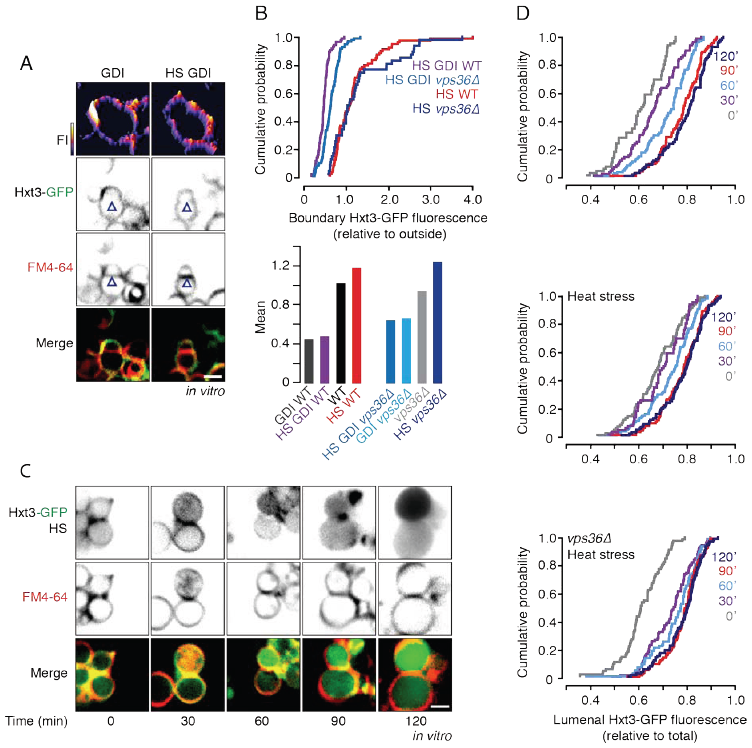
(**A**) Fluorescence micrographs of lysosomes isolated from wild type cells expressing Hxt3-GFP after 30 minutes of fusion in the presence of the fusion inhibitors rGdi1 and rGyp1-46 (GDI) alone or with heat stress (HS) treatment. (**B**) Cumulative probability plots (top) of Hxt3-GFP boundary fluorescence of lysosomes isolated from wild type (WT) or *vps36*∆ cells after fusion in the presence or absence of heat stress (HS) or fusion inhibitors (GDI). Mean values (below) are also shown and include values calculated from plots shown in **Figure 2D** (n ≥ 96). (**C**) Fluorescence micrographs of lysosomes isolated from wild type cells expressing Hxt3-GFP acquired over the course of the fusion reaction in presence of heat stress (HS). (**D**) Cumulative probability plots of GFP lumenal fluorescence measured within lysosomes isolated from WT or *vps36∆* cells after 0, 30, 60, 90 or 120 minutes of fusion in the presence of heat stress, as shown in **Figures 2E and S2C**. Isolated organelles were stained with FM4-64 to label vacuole membranes. Scale bars, 2 μm.

**Figure S3 related to Figure 3.**
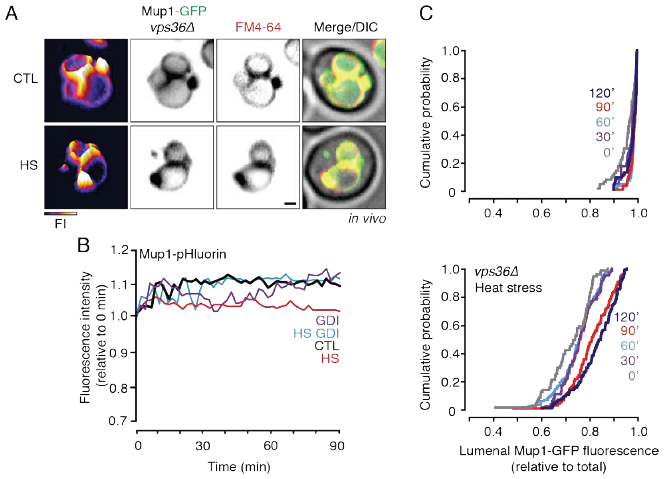
(**A**) Fluorescence and DIC micrographs of live *vps36*∆ cells expressing GFP-tagged Mup1 before (CTL) or after heat stress (HS). 3-dimensional fluorescence intensity (FI) plots of Mup1-GFP are shown. Isolated organelles were stained with FM4-64 to label vacuole membranes. Scale bars, 1 μm. (**B**) Fluorescence intensity of lysosomes isolated from wild type cells expressing Mup1-pHluorin during the *in vitro* fusion reaction under control conditions (CTL) or after heat stress (HS) in presence or absence of fusion inhibitors rGdi1 and rGyp1-46 (GDI). (**C**) Cumulative probability plots of Mup1-GFP lumenal fluorescence within lysosomes isolated from WT (top) or *vps36∆* cells treated with heat stress (bottom) after 0, 30, 60, 90 or 120 minutes of fusion, as shown in **Figure 3F**.

**Figure S4 related to Figure 4.**
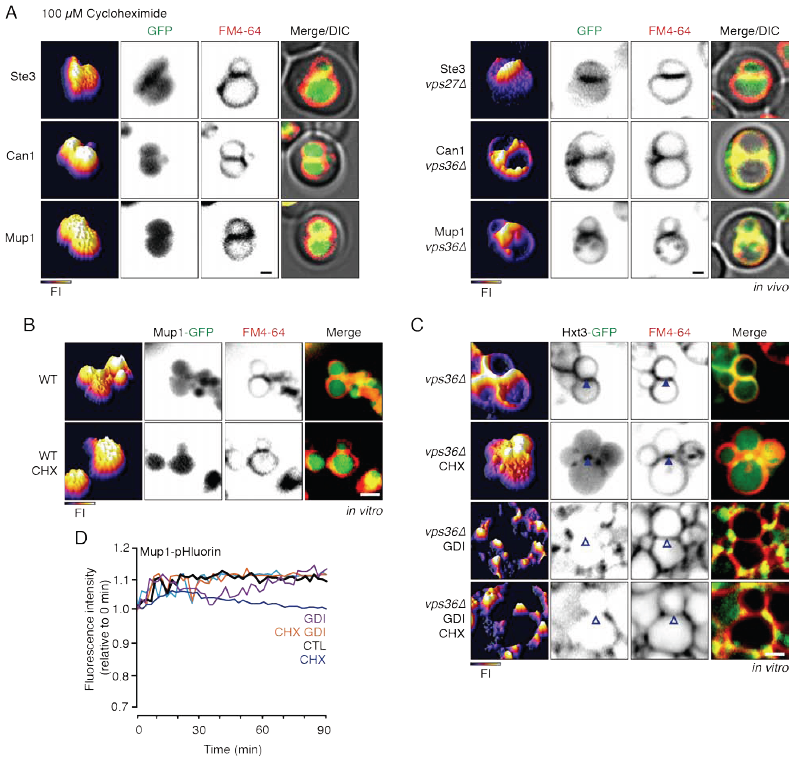
(**A**) Fluorescence and DIC micrographs of live wild type (left) and ESCRT deficient (*vps27*∆ or *vps36*∆; right) cells expressing GFP-tagged Ste3, Can1 or Mup1 after incubation with 100 μM cycloheximide for 70 minutes. (**B and C**) Fluorescence micrographs of lysosomes isolated from wild type Mup1-GFP cells after 15 minutes of fusion in the absence (WT) or presence of cycloheximide (CHX; **B**), or from lysosomes isolated from *vps36∆* cells expressing Hxt3-GFP in the absence or presence of cycloheximide (CHX) or rGdi1 and rGyp1-46 (GDI; **C**). 3-dimensional fluorescence intensity (FI) plots of Mup1-GFP or Hxt3-GFP are shown. Arrowheads indicate GFP-tagged protein enrichment (closed) or exclusion (open) within the boundary membrane. (**D**) Fluorescence of lysosomes isolated from wild type cells expressing Mup1-pHluorin measured during the *in vitro* fusion reaction under control conditions (CTL) or after cycloheximide (CHX) treatment in the presence or absence of the fusion inhibitors rGdi1 and rGyp1-46 (GDI). Cell or isolated organelles were stained with FM4-64 to label vacuole membranes. Scale bars, 1 μm *in vivo* and 2 μm *in vitro*.

**Table S1.** Yeast strains used in this study

| Strain | Genotype | Source |
| --- | --- | --- |
| SEY6210 | *MATα, leu1-3, 112 ura3-52 his3-200, trp1-901 lys2-801 suc2-D9* | *Huh et al., 2003* |
| BY4741 | *MATα his3-Δ1 leu2-Δ0 met15-Δ0 ura3-Δ0* | *Huh et al., 2003* |
| Fet5-GFP | BY4741, *Fet5-GFP::HIS3MX* | *Huh et al., 2003* |
| Mup1-GFP | SEY6210, *Mup1-GFP::KanMX* | *Prosser et al., 2011* |
| Mup1-GFP:*vps27*Δ | SEY6210, *Mup1-GFP::KanMX, vps27*Δ:*HIS3MX* | This study |
| Mup1-GFP:*vps36*Δ | SEY6210, *Mup1-GFP::KanMX, vps36*Δ:*HIS3MX* | This study |
| Mup1-pHluorin | SEY6210, *Mup1-pHluorin::KanMX* | *Prosser et al., 2011* |
| Mup1-pHluorin:*vps27*Δ | SEY6210, *Mup1-pHluorin::KanMX, vps27*Δ:*HIS3MX* | This study |
| Mup1-pHluorin:*vps36*Δ | SEY6210, *Mup1-pHluorin::KanMX, vps36*Δ:*HIS3MX* | This study |
| Ste3-GFP | SEY6210, *Ste3-GFP::KanMX* | *Prosser et al., 2011* |
| Ste3-GFP:*vps27*Δ | SEY6210, *Ste3-GFP::KanMX, vps27*Δ:*HIS3MX* | This study |
| Can1-GFP | BY4741, *Can1-GFP::HIS3MX* | *Huh et al., 2003* |
| Can1-GFP:*vps36*Δ | BY4741, *Can1-GFP::HIS3MX, vps36*Δ:*KanMX* | This study |
| Hxt3-GFP | BY4741, *Hxt3-GFP::His3MX* | *Huh et al., 2003* |
| Hxt3-GFP:*vps36*Δ | BY4741, *Hxt3-GFP::His3MX, vps36*Δ:*KanMX* | This study |
| Hxt3-GFP:*vps27*Δ | BY4741, *Hxt3-GFP::His3MX, vps27*Δ:*KanMX* | This study |
| Itr1-GFP | BY4741, *Itr1-GFP::His3MX* | *Huh et al., 2003* |
| Itr1-GFP:*vps27*Δ | BY4741, *Itr1-GFP::His3MX, vps27*Δ:*KanMX* | This study |
| Aqr1-GFP | BY4741, *Aqr1-GFP::His3MX* | *Huh et al., 2003* |
| Aqr1-GFP:*vps27*Δ | BY4741, *Aqr1-GFP::His3MX, vps27*Δ:*KanMX* | This study |

